# Repetitive mild concussion in subjects with a vulnerable cholinergic system: lasting cholinergic-attentional impairments in CHT^+/-^ mice

**DOI:** 10.1101/521484

**Authors:** Ajeesh Koshy Cherian, Natalie C. Tronson, Vinay Parikh, Aaron Kucinski, Randy D. Blakely, Martin Sarter

**Affiliations:** Department of Psychology and Neuroscience Program, University of Michigan, Ann Arbor, MI 48109, USA; Department of Psychology and Neuroscience Program, Temple University, Philadelphia, PA 19122, USA; FAU Brain Institute, Charles E. Schmidt College of Medicine, Florida Atlantic University, Jupiter, FL 33458, USA

**Author notes:** Correspondence: Martin Sarter, Department of Psychology and Neuroscience Program, University of Michigan, 530 Church Street, 4030 East Hall, Ann Arbor, MI 48109.

**Keywords:** Acetylcholine, Choline transporter, Concussion, Traumatic brain injury, Attention

## Abstract

Previous research emphasized the impact of traumatic brain injury on cholinergic systems and associated cognitive functions. Here we addressed the converse question: Because of the available evidence indicating cognitive and neuronal vulnerabilities in humans expressing low-capacity cholinergic systems or with declining cholinergic systems, do injuries cause more severe cognitive decline in such subjects, and what cholinergic mechanisms contribute to such a vulnerability? Using mice heterozygous for the choline transporter (CHT^+/-^ mice) as a model for a limited cholinergic capacity, we investigated the cognitive and neuronal consequences of repeated, mild concussion injuries (rmCc). Following five rmCc, and compared with WT mice, CHT^+/-^ mice exhibited severe and lasting impairments in sustained attention performance, consistent with effects of cholinergic losses on attention. However, rmCc did not affect the integrity of neuronal cell bodies and did not alter the density of cortical synapses. As a cellular mechanism potentially responsible for the attentional impairment in CHT^+/-^ mice, we found that rmCc nearly completely attenuated performance-associated, CHT-mediated choline transport. These results predict that subjects with an already vulnerable cholinergic system will experience severe and lasting cognitive-cholinergic effects following even relatively mild injuries. If confirmed in humans, such subjects may be excluded from, or receive special protection against, activities involving injury risk. Moreover, the treatment and long-term outcome of traumatic brain injuries may benefit from determining the status of cholinergic systems and associated cognitive functions.

---

Basal forebrain (BF) cholinergic projections to telencephalic regions mediate the integration of extero- and interoceptive cues into ongoing behavior and foster experience-based neuroplasticity (Ballinger, Ananth, Talmage, & Role, 2016; Conner, Chiba, & Tuszynski, 2005; Sarter, Albin, Kucinski, & Lustig, 2014). Among the behavioral and cognitive functions which depend on cholinergic activity, attentional performance, including the detection of cues (as defined by Posner, Snyder, & Davidson, 1980), and the capacity for attentional (or “top-down”) control, is closely linked to cholinergic signaling (Berry, Blakely, Sarter, & Lustig, 2015; Gritton et al., 2016; Howe et al., 2013; Howe et al., 2017; Kim, Muller, Bohnen, Sarter, & Lustig, 2017a; Parikh, Kozak, Martinez, & Sarter, 2007; Sarter, Lustig, Berry, et al., 2016; Sarter, Lustig, Blakely, & Koshy Cherian, 2016; Sarter, Lustig, Howe, Gritton, & Berry, 2014; Sarter & Paolone, 2011; St Peters, Demeter, Lustig, Bruno, & Sarter, 2011). As would be expected from such fundamental roles of cholinergic systems, cholinergic losses initiate and exacerbate cognitive-behavioral decline in patients with neurodegenerative disorders (Bohnen et al., 2003; Bohnen et al., 2009; Hampel et al., 2018; Kim et al., 2017a; Kim, Muller, Bohnen, Sarter, & Lustig, 2017b; Sarter, Albin, et al., 2014).

The exquisite vulnerability of cholinergic neurons to the long-term effects of traumatic brain injury has been extensively documented (e.g., Conner et al., 2005; Dixon, Bao, Long, & Hayes, 1996; Dixon, Hamm, Taft, & Hayes, 1994; Dixon et al., 1995; Salmond, Chatfield, Menon, Pickard, & Sahakian, 2005). Here we addressed the converse question: What are the cholinergic-attentional consequences of brain injury in subjects with an already vulnerable cholinergic system? This question arose in part from our research on human subjects expressing a low-capacity choline transporter (CHT) who exhibit attentional and affective vulnerabilities (Berry et al., 2015; Berry et al., 2014; English et al., 2009; Hahn et al., 2008; Sarter, Lustig, Blakely, et al., 2016), and on patients with Parkinson’s disease who, in addition to basal ganglia dopamine losses, exhibit cholinergic losses and attentional impairments which are correlated with their propensity for falls (Kim et al., 2017a, 2017b; Sarter, Albin, et al., 2014). Given these cholinergic-attentional risk factors, are these subjects also at a greater risk for developing severe impairments following traumatic brain injury?

We addressed this question by examining the attentional and cholinergic consequences of repeated mild injuries in mice heterozygous for the CHT (CHT^+/-^ mice). Compared to wild type (WT) mice, CHT^+/-^ mice model a cholinergic system that functions at WT levels at baseline but exhibits a limited capacity for sustaining elevated levels of cholinergic neurotransmission, such as during the performance of a Sustained Attention Task (SAT; Ferguson et al., 2004; Paolone, Angelakos, Meyer, Robinson, & Sarter, 2013; Parikh, St. Peters, Blakely, & Sarter, 2013). However, in the absence of additional challenges on performance or cholinergic neurotransmission, residual ACh release is sufficient to support basic SAT performance in these mice, similar to preserved performance but vulnerability to distractors in humans expressing a low-capacity CHT variant. Thus, CHT^+/-^ mice serve as a useful model to study interactions with an expectedly detrimental manipulation of cholinergic function.

The literature on the effects of traumatic brain injury is complex, in part because seemingly subtle variations in the apparatus and methods used to administer injuries generate substantial differences in primary and secondary injuries (Jassam, Izzy, Whalen, McGavern, & El Khoury, 2017; Xiong, Mahmood, & Chopp, 2013). As has been widely discussed, repeated mild concussion injuries (rmCc) are common brain injuries (e.g., Collins et al., 1999; Corrigan, Selassie, & Orman, 2010). Therefore, we investigated the relative vulnerability of CHT^+/-^ mice to rmCc. We used a modified version of the impact apparatus and method described by Kane et al. (2012; see Methods for details) to administer repeated yet relatively even milder concussion injuries. rmCC produced lasting attentional impairments in CHT^+/-^ mice which were associated with a near complete silencing of CHT function.

## Materials and Methods

### Subjects and housing conditions

Male and female mice (N = 76; see Results for the number of mice per genotype, sex and experimental condition), heterozygous for the choline high-affinity transporter (CHT^+/−^) were originally obtained from Vanderbilt University (Ferguson et al., 2004), maintained on a C57BL/6 genetic background for over seven generations and continued to be bred on this background at the University of Michigan. Mice were at least 12 weeks of age and weighed 20-30 g at the beginning experiments.

Mice were genotyped by a commercial vendor (Transnetyx, Cordova, TN). CHT^+/−^ (n = 42) and wild type mice (WT; n = 40) were housed individually in a temperature (23°C) and humidity-controlled (45%) environment, and kept on a 12:12 light/dark cycle (lights on at 7:30 a.m.). Training and testing on the Sustained Attention task (SAT) took place during the light phase. We previously demonstrated that SAT practice evokes a diurnal activity pattern in rats (Gritton, Kantorowski, Sarter, & Lee, 2012; Gritton, Sutton, Martinez, Sarter, & Lee, 2009; Paolone, Lee, & Sarter, 2012), suggesting that mice likewise practiced the SAT during their active period. Mice not performing the SAT had *ad libitum* access to food and water. SAT-performing mice were gradually water-deprived over a five-day period prior to the onset of training. Water was then restricted to a 4-min period following each training session. During SAT sessions, correct responses were rewarded using sweetened water (0.2% saccharin; 6 μL per reward; total average session delivery: 0.45 mL). On days without SAT practice, water access in the animals’ home cages was increased to 10 min. Food (Rodent Chow, Harlan Teklad, Madison, WI) was available *ad libitum*. During SAT training, animals’ body weights remained stable at 90-100% of their *ad libitum* body weights (weighed twice weekly). SAT performing mice were trained 5-6 days per week. All procedures were conducted in adherence with protocols approved by the University of Michigan Institutional Laboratory Animal Care and Use Committee (ILACUC} and conducted in laboratories accredited by the Association for Assessment and Accreditation of Laboratory Animal Care (AAALAC).

### SAT apparatus, task acquisition, and performance measures

Behavioral training and testing took place using twelve modified operant chambers (24.10 cm L × 20.00 cm W × 29.50 cm H; MedAssociates, Inc., St. Albans, VT), situated inside sound-attenuated chambers fabricated at the University of Michigan. Each operant chamber contained an intelligence panel equipped with two panel lights (2.8 W), two retractable “Michigan Controlled Access Response Ports” (MICARPs; St Peters, Cherian, Bradshaw, & Sarter, 2011), and a liquid dispenser (6 μL of 0.2% saccharin in de-ionized water per delivery; Fig. 1a). Stimuli and response recordings were implemented using SmartCtrl Package 8-In/16-Out with additional interfacing by MED-PC for Windows (Med Associates, Inc., St. Albans, VT) and custom programming. The MICARPs were located laterally on either side of the liquid dispenser, and the two panel lights were located directly above the liquid dispenser. The MICARPs and the liquid dispenser each contained perpendicular infrared photo-beams to detect nose-poke responses. A house light (2.8 W) was located at the top of the rear wall. Each sound-attenuating chamber was also equipped with a ventilation fan, a video camera, and 4 red LEDs (JameCo P/N 333489; Jameco Valuepro, Belmont, CA).

**Figure 1.**
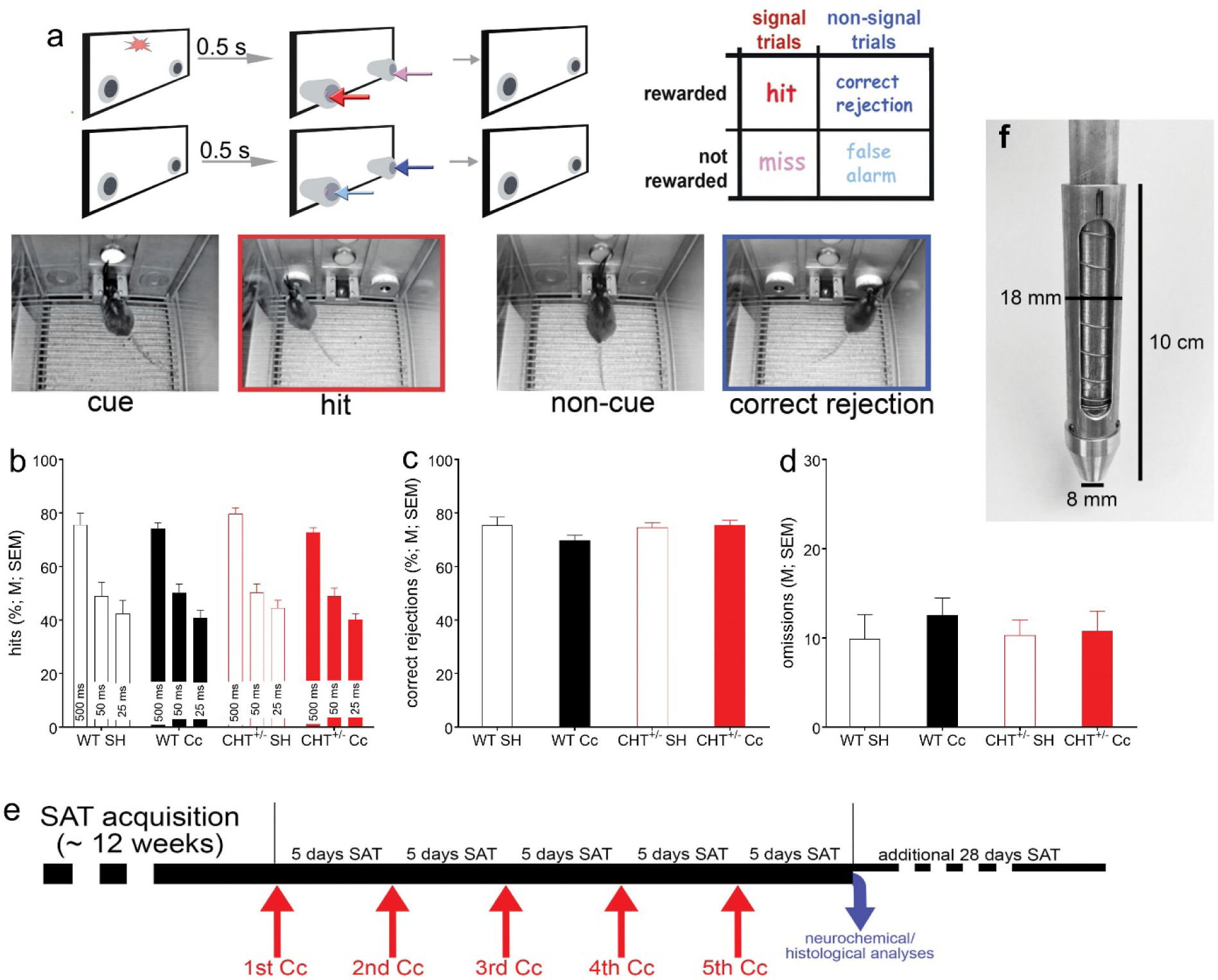
Illustration of the Sustained Attention Task (SAT) and pre-rmCc performance of WT and CHT^+/-^ mice. <u>a</u>: The SAT consisted of a randomized sequence of signal or cue (illumination of the center panel light; see left photograph) and non-signal (or non-cue) events, followed by extension of the two response ports. Hits and correct rejections, but not misses and false alarms, were rewarded. Note the matching colors of the arrows pointing to the response ports for the four response categories and the arrows in the outcome matrix on the top right, indicating that, in cued trials, hits (red arrows) are rewarded and in non-cued trials, correct rejections were rewarded (dark blue). Conversely, misses (pink arrows) and false alarms (light blue arrows) were not rewarded and triggered an intertrial interval. The photographs depict a cue event (left), followed by a hit (2^nd^ from the left, red frame, matching the arrow color), and non-cued trial (third from left) followed by a correct rejection (dark blue frame). <u>b-d</u>: Prior to the administration of rmCc, SAT performance did not differ between genotypes and mice assigned to receive either SH or rmCc (for ANOVAs see Results; WT SH: n = 6; WT rmCc: n = 15; CHT^+/-^ SH: n = 9; CHT^+/-^ rmCc: n = 14). Neither the relative number of hits to 500, 50, and 25-ms signals (<u>b</u>), the relative number of correct rejections (<u>c</u>), nor the number of omissions (out of a total of 170 trials /session; <u>d</u>) differed between the genotypes and assignment to SH or rmCc. e illustrates the timeline of the main experiment. Following SAT acquisition, five rmCc events were separated individually by 5 days of SAT practice. After the final 5 days of practice, after the 5^th^ event, the brains of the majority of mice were harvested for neurochemical and histological analyses while a small proportion of rmCc-treated mice (n = 4 per genotype) continued daily SAT practice for an additional 28 sessions to determine lasting effects. <u>f</u> shows the transfer ram of the rmCc apparatus. The addition of this transfer ram constituted the main modification of the apparatus previously used by Kane et al. (2012) to administer repeated, relatively mild impacts in mice. This transfer ram was equipped with an internal compression ring to further limit the impact of the weight drop.

The task, training procedures, and validity of the SAT performance measures in terms of indicating sustained attention in humans, rats, and mice were previously described (Demeter, Sarter, & Lustig, 2008; Paolone, Mallory, et al., 2013; St Peters, Cherian, et al., 2011). Briefly, animals were initially shaped to reach the water dispenser using a modified fixed ratio-1 (FR-1) reward schedule. After obtaining ≥100 water rewards within the 40-min training session for two consecutive days, animals began SAT training. First, mice were familiarized with moving MICARPS. Both MICARPS extended until a nose-poke triggered retraction. Mice were required to generate 90 rewards/session and retrieve at least 80% of these rewards within 1.5 s after the nose poke for two consecutive sessions before moving to the next training stage. During all subsequent training stages, mice were required to distinguish between signal (illumination of the central panel light) and non-signal (no illumination) events by responding to the respective MICARP. The house lights of the cage were kept off until the last stage of training, to increase the relative saliency of the signal and facilitate the learning of the task rules. On signal trials, a left nose-poke was reinforced and termed a “hit.” On non-signal trials, a right nose-poke was reinforced and termed a “correct rejection” (for an illustration of trial types see Fig. 1a). Half of the animals were trained following the reverse rules. Incorrect responses to signal trials (“misses”) and non-signal trials (“false alarms”) triggered an inter-trial interval (12 ± 3 s) but had no further scheduled consequences. An omission was recorded if the animal failed to nose-poke during the lever extension following an event. Signal and non-signal trials were presented in a pseudo-randomized order.

In the first training stage, the signal light remained illuminated for 500 ms on signal trials. MICARPs were extended 1 s after the signal onset and remained extended for 4 s or until a nose poke triggered a retraction. During non-signal trials, the MICARPs were likewise extended for 4 s or until a nose poke was initiated. Incorrect responses were followed with up to three correction trials, during which the previous trial was repeated. If the animal failed to respond correctly to all three correction trials, a forced trial was initiated, during which only the MICARP corresponding to the correct response extended. After reaching stable performance, defined by at least three consecutive days of obtaining >60% hits and correct rejections with <30% omissions, signal length and time delay before nose-poke extension were reduced (1 sec and 0.5 s, respectively). When animals again reached stable performance, multiple signal durations were introduced (500, 50, or 25 ms). The session length was 40 min, allowing for *post hoc* analysis of performance over five 8-min blocks. Pseudo-randomization was designed to ensure equal numbers of signal and non-signal trials as well as trials with 500, 50 or 25 ms signals (approximately 160 trials/session). Forced correction trials were eliminated at this point. After reaching stable performance, animals moved to the final stage training and the actual SAT task. During this stage, the operant chamber house light was illuminated, requiring animals to sustain orientation towards the signal panel. Criterion performance for undergoing rmCc was defined as an average of >60% hits to 500 ms trials and correct rejections, <30% omissions for five consecutive days. Hits, misses, correct rejections, false alarms, and omissions were recorded for each SAT session. The relative number of hits (hits/hits + misses) was calculated for each signal length, as well as the relative number of correct rejections (correct rejections/correct rejections + false alarms). Measures of performance were calculated separately for each of the five, 8-min task blocks.

### rmCc: apparatus and procedure

The majority of weight drop impactors described in the literature were designed to produce relatively severe injuries, evidenced by macroanatomical lesions (e.g., Marmarou et al., 1994). Other impactors were used to produce relatively milder and more diffuse injuries but associated procedures limited secondary impacts associated with abrupt rotational head movements which often characterize the impacts suffered by athletes or military personnel (e.g., Milman, Rosenberg, Weizman, & Pick, 2005; Xiong et al., 2013; Zohar et al., 2003). The rmCC apparatus used in the present study was adopted from Kane et al. (2012; see their Figure 1) but modified to produce a relatively even less severe impact. The apparatus used the energy from a falling weight (190 g; drop height: 50 cm) to induce a rapid acceleration onto the head of an unrestrained mouse. The rig consisted of a vertical rod supported from above, a sliding platform and, importantly, a transfer ram equipped with a weight transfer ram with an internal compression spring to absorb a portion of the impact force (Fig. 1f). The transfer ram weighted 20 g, the compression spring (Hillman Group, Cincinnati, OH; Item #540281, #94 compression spring) had a free length of 6.35 cm long, 1.35 cm when fully compressed, 1.42 cm in outer diameter, and it had a 0.66 mm wire diameter. The concussion platform consisted of a Plexiglas chamber (15 cm length X 10 cm width X 10 cm height) with a replaceable slit aluminum foil bed at the top and a foam cushion at the base. Because the impact of the weight interacted with a mouse falling though an aluminum foil “hammock” (below), an estimate of the force reduction afforded by the spring inside the transfer ram (which pushed back on the falling weight and thus effectively reduced its weight), assumed a stationary mouse (25 g) and suggested 1.96 N without and 1.22 N with the spring (a reduction of 38%).

For administering an impact mice were lightly anesthetized with isoflurane and placed chest down on the bed of aluminum foil. The vertical rod was adjusted so that the transfer ram rested on the animal’s head between the ears. The sliding platform was loaded, raised to a pre-determined height, and was then released. As the sliding platform hit the bottom of the rod, the transfer ram moved rapidly, imparting a rapid acceleration to the head of the animal. Immediately following the impact, the animal fell freely through the slit aluminum foil onto a foam pad, placed 10 cm below. This arrangement of the apparatus avoided any crushing type of injury as the head of the animal was not restrained, and it ensured a mild concussive injury produced by the high velocity impact and rapid acceleration of the free moving head and body. Thereafter the mice were immediately transferred to their home cage. The total number of impacts (five) that were administered in the present experiment was based on the results described by Kane et al. (2012), indicating that 5 impacts (over 5 days) did not disrupt vital functions, the integrity of the skull and brain, and motor performance. Furthermore, our pilot studies indicated that administering 5 impacts over 5 weeks, using our modified apparatus, spared the SAT performance of WT mice but impaired the hit rates of CHT^+/-^ mice. Thus, our modified apparatus and administration regimen were designed to yield a robust performance contrast between the genotypes.

### Choline uptake assay

Performance-associated CHT capacity was measured in cortical synaptosomes isolated immediately after the final SAT session (7 days after the 5^th^ rmCc; for a schematic timeline see Fig. 1e) and from a subset of mice from all groups (N = 16; 4 mice per genotype and condition). Mice were decapitated under urethane anesthesia. Synaptosomes were prepared from isolated tissue as described earlier (Parikh et al., 2013). Briefly, right and left frontal cortices were pooled and homogenized in ice-cold 0.32 M sucrose. The homogenate was centrifuged at 1000 X g for 4 min at 4^0^C to remove cellular debris. A synaptosomal pellet was obtained by spinning the resultant supernatant at 12,500 X g for 10 min. Aliquots (50 µL) of crude synaptosomes were incubated with 100 µL [^3^H]-choline (0.02-6.0 µM) in Krebs bicarbonate buffer in the presence and absence of 10 µM hemicholinium-3 (HC-3) for 5 min at 37^0^C. Transport assays were terminated by transferring the tubes to an ice bath followed by rapid filtration using a cell harvester (Brandel Inc., Gaithersburg, MD). Accumulated radioactivity was determined using a liquid scintillation counter. Protein concentration was measured using Pierce BCA protein assay kit (Thermo Fisher scientific Inc., Rockford, IL). CHT-mediated choline uptake was determined as total choline uptake minus the uptake in the presence of hemicholinium-3 and maximum transporter velocity (Vmax) and the affinity for choline (Km) were determined.

### Western Blotting

Non-performing WT and CHT+/− mice (N = 16; n = 3-5 per group) underwent a total of five rmCc or sham procedures, following the same timeline for rmCc delivery as was applied to the SAT-performing animals (see Fig. 1e). Seven days after the final impact, mice were decapitated under urethane anesthesia. Brains were removed immediately and frontal cortical tissues were dissected on an ice-cold petri dish. Isolated tissues pooled from both hemispheres were homogenized in ice-cold 0.32M sucrose. The homogenate was centrifuged at 1000 X g for 4 min at 4°C to remove cellular debris. Supernatants were centrifuged at 12,000 X g for 15 min at 4°C to obtain the crude synaptosomal pellet. Synaptosomes were resuspended in the lysis buffer containing 100 mM Tris-HCl, 50 mM NaCl, 1 mM EDTA, 1 mM EGTA, 1% SDS, 1% Triton X-100 and protease inhibitor cocktail. Protein concentrations were determined by using a modified Lowry Protein Assay (Pierce, Rockford, IL). CHT immunoblotting was conducted as described in our previous studies (Apparsundaram, Martinez, Parikh, Kozak, & Sarter, 2005; Parikh, Apparsundaram, Kozak, Richards, & Sarter, 2006; Parikh et al., 2013). Briefly, samples for SDS-PAGE were prepared in Laemmli buffer by heating at 80^0^C for 20 min prior to loading. Proteins (25 µg/sample) were separated on 4-15% Tris HCl polyacrylamide gels and transferred on PVDF membranes. A mouse anti-CHT monoclonal antibody (clone 62-2E8; EMD Millipore, Temecula, CA) was used at 1:1000 dilution for the immunodetection of CHT bands. The blots were exposed to a peroxidase-conjugated anti-mouse secondary antibody and ECL Advance chemiluminescent substrate (GE Healthcare, Piscataway, NJ). The resulting chemilluminescent signal was acquired with a Molecular Imager Chemidoc EQ system (Bio-Rad, Hercules, CA). All membranes were stripped in Restore Plus buffer (Pierce) and re-probed with rabbit anti-synaptophysin antibody (Sigma-Aldrich) to detect synaptophysin that served as a general marker for presynaptic terminal proteins. Densitometric analysis of immunoreactive bands was performed by calculating the integrated pixel densities using NIH ImageJ software. To accommodate for any differences in sample loading across different samples, the density of CHT- and synaptophysin-immunoreactive bands was normalized to the levels of β-tubulin-immunoreactive bands for each sample analyzed.

### Histological analyses

Following the last five days of SAT practice (5 days after the 5^th^ rmCc administration; Fig. 1e), the brains from a subgroup of animals were harvested for histological analyses. Animals were given an overdose of sodium pentobarbital and perfused with 0.1% phosphate buffer solution (PBS) followed by 4% paraformaldehyde in 0.15 M phosphate buffer and 15% saturated picric acid, pH 7.4. Brains were post-fixed for 4 h and were then rinsed in 0.1 M PBS, followed by storing in 30% sucrose PBS solution until they sank. Coronal sections (35 μm) were sliced using a freezing microtome (CM 2000R; Leica) and stored in antifreeze solution until further processing. For assessing macroanatomical damage and gliosis, Nissl-staining was performed on sections mounted on gelatin coated slides after allowing them to dry completely.

### Statistical analyses

SAT performance measures (hits, correct rejections and omissions) were compared between genotypes (WT or CHT^+/-^) and treatments (sham or rmCc) using repeated-measures or one- or two-way ANOVAs. Sex was a factor in all analyses. The analysis of hits also included the within-subject factor signal duration (500, 50, and 25 ms). Following significant main effects, *post hoc* multiple comparisons were conducted using the least significant difference (LSD) test or Mann-Whitney U Test. Statistical analyses were performed using SPSS for Windows (version 17.0). The Kruskal Wallis test was used to compare CHT-mediate choline uptake and CHT protein levels between experimental groups. In cases of violation of the sphericity assumption, Huyhn–Feldt-corrected *F*-values, along with uncorrected degrees of freedom, are given. Alpha was set at 0.05 or 0.05/3 when designated. Exact *P* values are reported as recommended previously (Greenwald, Gonzalez, Harris, & Guthrie, 1996). Variances are reported and illustrated as standard error of the mean (SEM). Effect sizes for selected effects are reported using Cohen’s *d* (Cohen, 1988).

## Results

### rmCc-induced severe and lasting attentional impairments in CHT^+/-^ mice

WT (n = 21, 12 females) and CHT^+/-^ mice (n = 23, 15 females) were trained in the SAT (Fig. 1a) to performance criterion (>60% hits to 500 ms signals, >60% correct rejections, and <30% omissions for five consecutive days/sessions). The number of sessions to reach this criterion did not differ between WT and CHT^+/-^ mice (t(42) = 0.80, *P* = 0.43; WT: 98.57 ± 12.98 sessions; CHT^+/-^: 86.74 ± 7.27 sessions).

Prior to the administration of rmCc, mice were assigned to the four experimental groups (WT sham (SH): n = 6 (3 females); WT rmCc: n = 15 (9 females); CHT^+/-^ SH: n = 9 (6 females); CHT^+/-^ rmCc: n = 14 (9 females)). SAT performance data obtained from the last 3 sessions prior to the first Cc administration confirmed that the relative number of hits varied by signal duration (F(2,72) = 310.27, *P* < 0.0001). As was expected based on prior evidence showing that basic SAT performance – in the absence of distractor challenges – does not differ between the genotypes (Paolone, Mallory, et al., 2013), presurgery SAT performance did not differ between the groups (main effect of genotype, (future) rmCc, and 3-way interactions involving the factor sex: all F < 1.56, all *P* > 0.21; main effect of sex: F(1,36) = 2.64, *P* = 0.11). Likewise, there were no effects of genotype, (future) rmCc, or sex and no interactions between these factors on the relative number of correct rejections (all F < 2.51, all *P* > 0.12; Fig. 1c) and the number of omitted trials (all F < 0.29, all *P* > 0.60; Fig. 1d). These findings is consistent with results from prior experiments indicating that, in the absence of behavioral or pharmacological challenges of SAT performance or cholinergic neurotransmission, the residual capacity of the cholinergic system of CHT^+/-^ mice is sufficient to support SAT performance (Paolone, Mallory, et al., 2013; Parikh et al., 2013).

Following SAT acquisition, mice were subjected to SH or rmCc using a slightly modified version of the method described by Kane et al. (2012; see their Figure 1 for an illustration of the suspended placement of mice, allowing unrestrained head acceleration upon impact). We modified this device to further minimize the effects of rmCc on the attentional performance of WT mice (see Methods for details). Mice were subjected to a total of five rmCc events over 35 days (see Fig. 1e for an illustration of the timeline). The mice practiced the SAT each day except on days when Cc was administered.

The analysis on the effects of genotype (WT, CHT^+/-^), Cc (SH, Cc), and rmCc event number (1-5) was conducted separately for hits to each of the three signal durations (with α=0.05/3). In the analysis of hits to longest signals, this 3-way ANOVA indicated a significant main effect of rmCc (F(1,36) = 13.20, *P* = 0.001) that interacted with genotype (F(1,36) = 15.18, *P* = 0.001; Fig. 2a,b). This result reflected that rmCc decreased hit rates in CHT^+/-^ but not in WT mice (Fig. 2a). After the 5^th^ rmCc event, hits to longest signals remained at pre-Cc levels in WT mice but decreased to about 50% in CHT^+/-^ mice (Cohen’s *d* = 1.80; Fig. 2b). The effects of rmCc and genotype on hits to shorter (50 ms and 25 ms) signals were not significant (all main effects and interactions: all F < 2.29, all P > 0.06), reflecting that hit rates to these signals were already relatively low prior to Cc administration (see also Fig. 1b).

**Figure 2.**
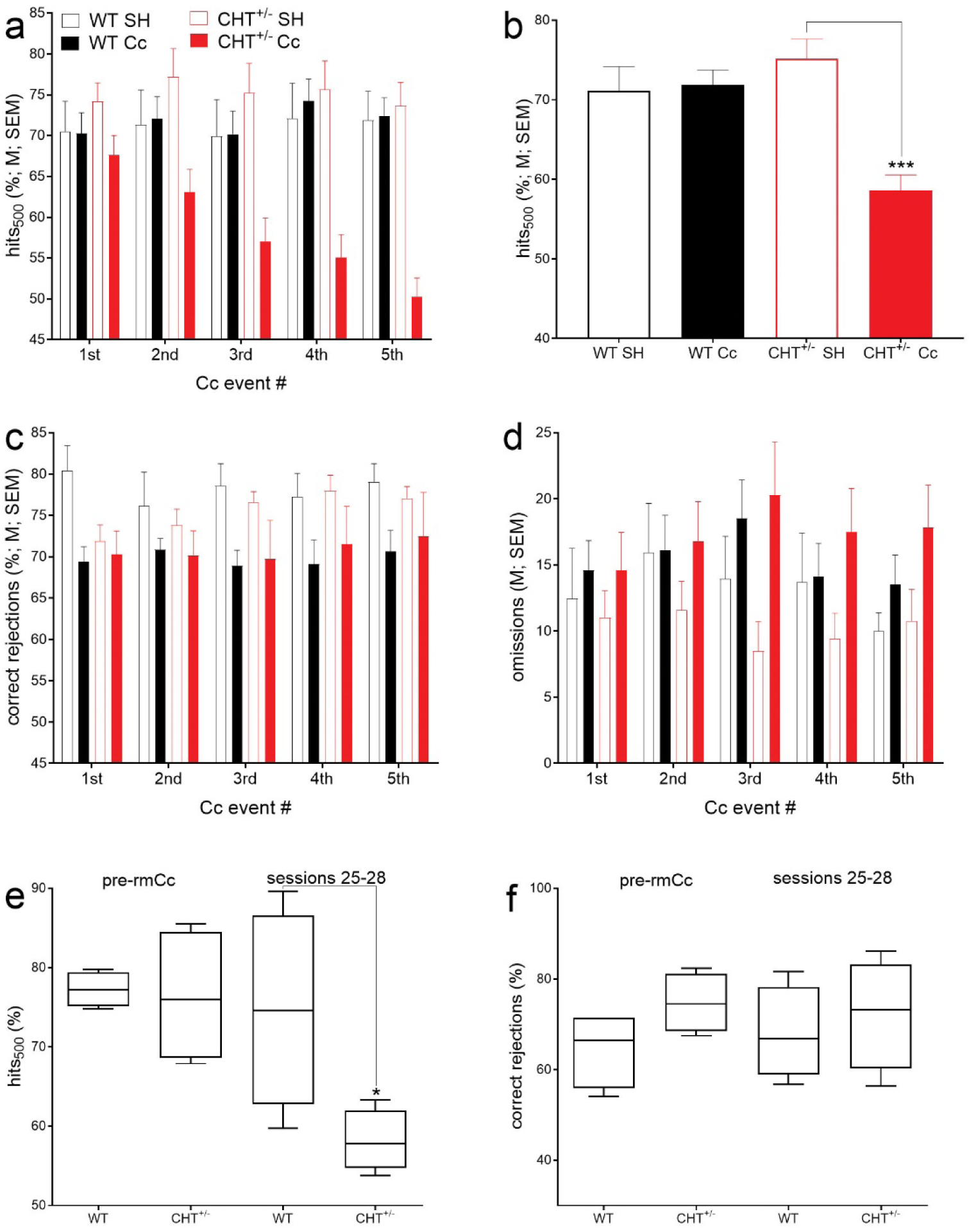
rmCc decreased hits to longest signals in CHT^+/-^ but not WT mice (<u>a</u>; 6-15 mice per group; note that Fig. 2a does not indicate multiple comparisons because the factor rmCc *number* (1-5) did not interact significantly with the effects of genotype). Concerning hits to longest signals, a significant interaction between the effects of genotype and rmCc (ANOVA in Results) reflected that, following rmCc, hit rates in CHT^+/-^ mice decreased to about 50% but remained at control levels in WT mice (<u>b</u>; ***, *P* < 0.001). There were no significant effects on hits to shorter signals, likely due to relatively low baseline hit rates to 50- and 25-ms signals (Fig. 1b). rmCc (main effect) significantly reduced correct rejections by about 7% in both genotypes (<u>c</u>). The number of omissions was not significantly affected by genotype or rm-TBI (<u>d</u>). A small proportion of rmCc-treated mice (n = 4/genotype) continued practicing the SAT for 28 days following the final impact. Continued SAT practice did not abolish the rmCc-induced decrease in hits in CHT+/-mice (<u>e</u>; boxes extend from the 25^th^ to 75 percentiles and whiskers show the range of data; medians are also shown). Correct rejections (<u>f</u>) did no longer differ between the genotypes during sessions 25-28.

Independent from genotype, the relative number of correct rejections were significantly decreased by rmCc (main effect of rmCc: F(1,36) = 6.36, *P* = 0.02); effects of genotype, sex and interactions: all F < 2.10, all *P* > 0.16; SH (M, SEM): 76.87 ± 2.16%; rmCc: 70.32 ± 1.53%; Fig. 2c). As detailed below, the relatively small decrease in correct rejections did not persist in the long-term, and may have reflected a non-cholinergically mediated decrement in instrumental performance (Burk & Sarter, 2001). The number of omitted trials was not affected by any factor (all F < 3.45, all P > 0.07).

#### Persistent decrease in hits

As the brains of the mice from this experiment were harvested for neurochemical and histological analyses following the final five days of SAT performance after the 5^th^ rmCc event, the persistence of the impairments in hits over longer periods following the 5^th^ Cc event was determined in a replication sample (n = 4 per genotype). These mice continued practicing the SAT for 28 sessions following the 5^th^ rmCc impact (see timeline in Fig. 1e). Compared WT-rmCc mice, continued daily SAT practice did not restore the rmCc-induced decrease of hit rates in CHT^+/-^ mice (U = 1, *P* = 0.04; Fig. 2e). The relative number of correct rejections (Fig. 2f) and the number of omission (not shown) did not differ between the genotypes during SAT sessions 25-28 (both *P* > 0.08).

### rmCc-induced decrease in CHT capacity

Immediately following completion of the final (5^th^) SAT session following the 5^th^ rmCc impact (Fig. 1e), brains were harvested for neurochemical and histological analysis. Our initial focus on CHT capacity and the integrity of cholinergic terminals (below) was based on two reasons. First, we previously demonstrated that the cholinergic neurons of CHT^+/-^ mice cannot support sustained increases in cholinergic neurotransmission (Paolone, Mallory, et al., 2013; Parikh et al., 2013). Because of prior evidence indicating that brain injuries disrupt cholinergic function (Dixon et al., 1996), we hypothesized that rmCc may further reduce CHT capacity in CHT^+/-^ mice and thereby permanently impair their SAT performance.

CHT capacity was determined in cortical synaptosomal preparations from 4 mice per genotype and condition which were harvested following the final SAT session. Saturation kinetic analysis (Fig. 3a) did not indicate effects of phenotype or rmCc on the affinity of choline to the CHT (median Km value across all 4 groups: 2.16 µM; 25^th^/75^th^ percentile: 1.39/3.86 µM; Kruskal Wallis test; H(3) = 2.74, *P* = 0.43). However, the capacity of the CHT, as indicated by maximal velocity (Vmax), differed significantly among the 4 groups (H(3) = 13.06 *P* < 0.0001, with rank sums of 58 for WT SH, 30 for WT rmCc, 38 for CHT^+/-^ SH and 10 for CHT^+/-^ rmCc; Fig. 3b). *Post hoc* comparisons between the groups indicated that WT Cc and both CHT^+/-^ groups had Vmax scores that were lower than those obtained from WT SH mice, that rmCc decreased Vmax scores in both genotypes, and that CHT^+/-^ rmCc mice had lower Vmax scores than WT rmCc mice (all U = 0, all *P* = 0.028).

**Figure 3.**
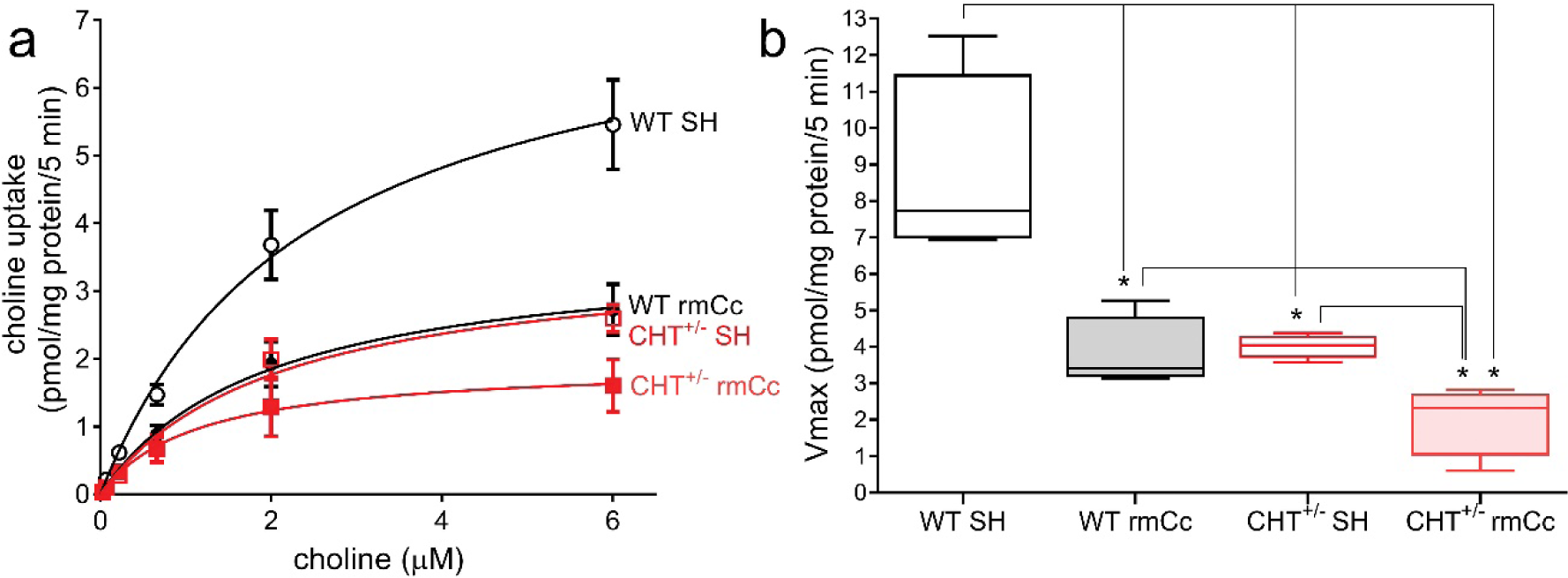
Cortical synaptosomal preparations were generated from mice which completed the last SAT practice session after the 5^th^ rmCc or SH event (n = 4 per genotype and treatment). Saturation kinetics analysis of hemicholinium (HC-3)-sensitive choline uptake (<u>a</u>) indicated significant group differences in Vmax (<u>b</u>) but not Km (not shown). <u>b</u> depicts boxes and whiskers (boxes depict the 25^th^ and 75 percentile, whiskers indicate the lowest and highest value) of Vmax scores. The results from a Kruskal Wallis test indicated a highly significant difference between the groups (see Results) that was followed up by non-parametric multiple comparisons shown here (*, *P* < 0.05). Note that previous studies demonstrated similar basal; (unstimulated) Vmax values in WT and CHT^+/-^ mice. Thus, the relatively lower Vmax values in CHT^+/-^ SH mice, when compared with WT SH mice, reflect SAT performance-induced stimulation of cholinergic activity, thereby revealing the limited outward trafficking of the CHT in CHT^+/-^ mice (Parikh et al., 2013). rmCc significantly reduced CHT capacity in WT and CHT^+/-^ mice, with the latter mice showing significantly lower Vmax values than the other three groups.

### rmCc reduced cortical CHT protein in WT mice but not the density of cholinergic terminals

Total CHT protein levels in the brain of CHT^+/-^ mice were expected to be approximately half of the levels measured in WT mice (Ferguson et al., 2004; Parikh et al., 2013). We measured total CHT levels in cortical synaptosomes, obtained from 3-5 mice per genotype and condition, and found a significant difference among the four groups (Kruskal Wallis test; H(3) = 9.11, *P* = 0.01; Figs. 4a,b). The effects of genotype failed to reach significance (U = 0, *P* = 0.057); however, median CHT values (normalized density) precisely reflected the expected impact of CHT heterozygosity (36.87 versus 18.54; Fig. 4b). rmCc significantly reduced CHT protein levels in WT mice (U = 0, *P* = 0.02; Fig. 4b) but had no further effect on CHT levels in CHT^+/-^ mice (U = 4, *P* = 0.62).

**Figure 4.**
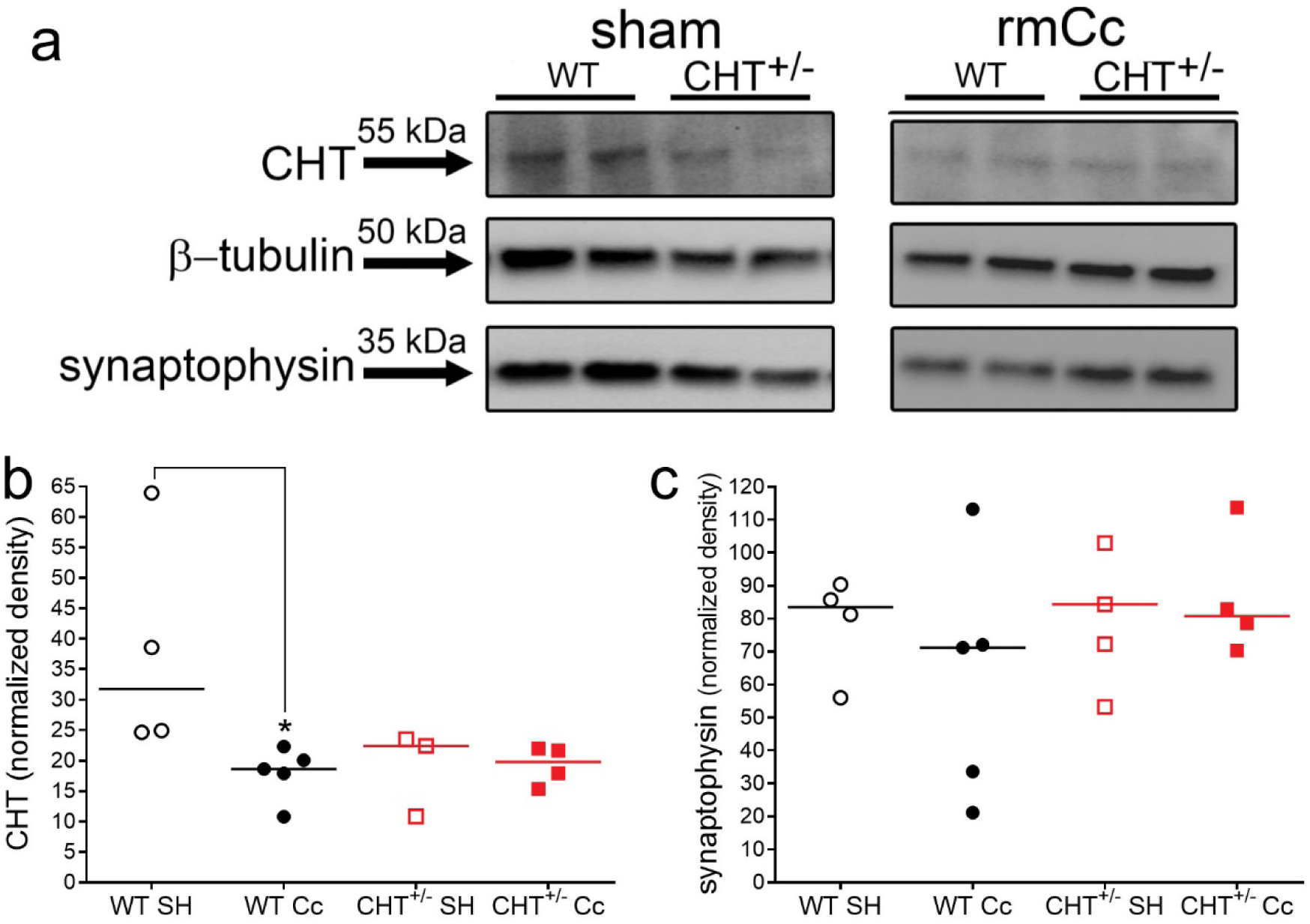
Immunoblot analysis of cortical CHT and synaptophysin protein levels in WT and CHT^+/-^ mice following SH or rmCc. <u>a</u> Immunoblots showed the expectedly reduced total CHT levels in CHT^+/-^ mice when compared to levels in WT SH mice (the two bands per group and condition indicate data from two mice each). Furthermore, the rmCc-induced decrease in CHT levels in WT mice is apparent. These examples also indicate that synaptophysin levels did not differ between the phenotypes and SH versus rmCc. <u>b</u> and <u>c</u> depict normalized, integrated CHT and synaptophysin levels, respectively (*, *P* < 0.05, based on Kruskal Wallis test and multiple non-parametric comparisons; the genotype effect in SH mice failed to reach significance; *P* = 0.057).

Because rmCc-induced decreased in hits mirror the effects of the removal of basal forebrain cholinergic neurons (e.g., McGaughy, Kaiser, & Sarter, 1996), we determined whether rmCc decreased the density of cortical terminals. Western blot analysis of the vesicular membrane protein synaptophysin (Tarsa & Goda, 2002; Thiel, 1993; Walaas, Jahn, & Greengard, 1988) did not indicate effects of rmCc or genotype on terminal density (*P* = 0.52; Figs. 4a,c). These data reject the possibility that rmCc resulted in the loss of cholinergic terminals and that such loss may explain the loss of rmCc-induced CHT capacity (Fig. 3).

### Histological analyses

Consistent with prior findings using this rmCc method (Kane et al., 2012), inspection of Nissl-stained brain sections did not indicate loss of cortical matter or disruption of the integrity of cortex and hippocampus (Fig. 5a-d). Thus, both synaptophysin protein levels and Nissl-stained sections indicated that absence of overt tissue damage resulting from rmCC.

**Figure 5.**
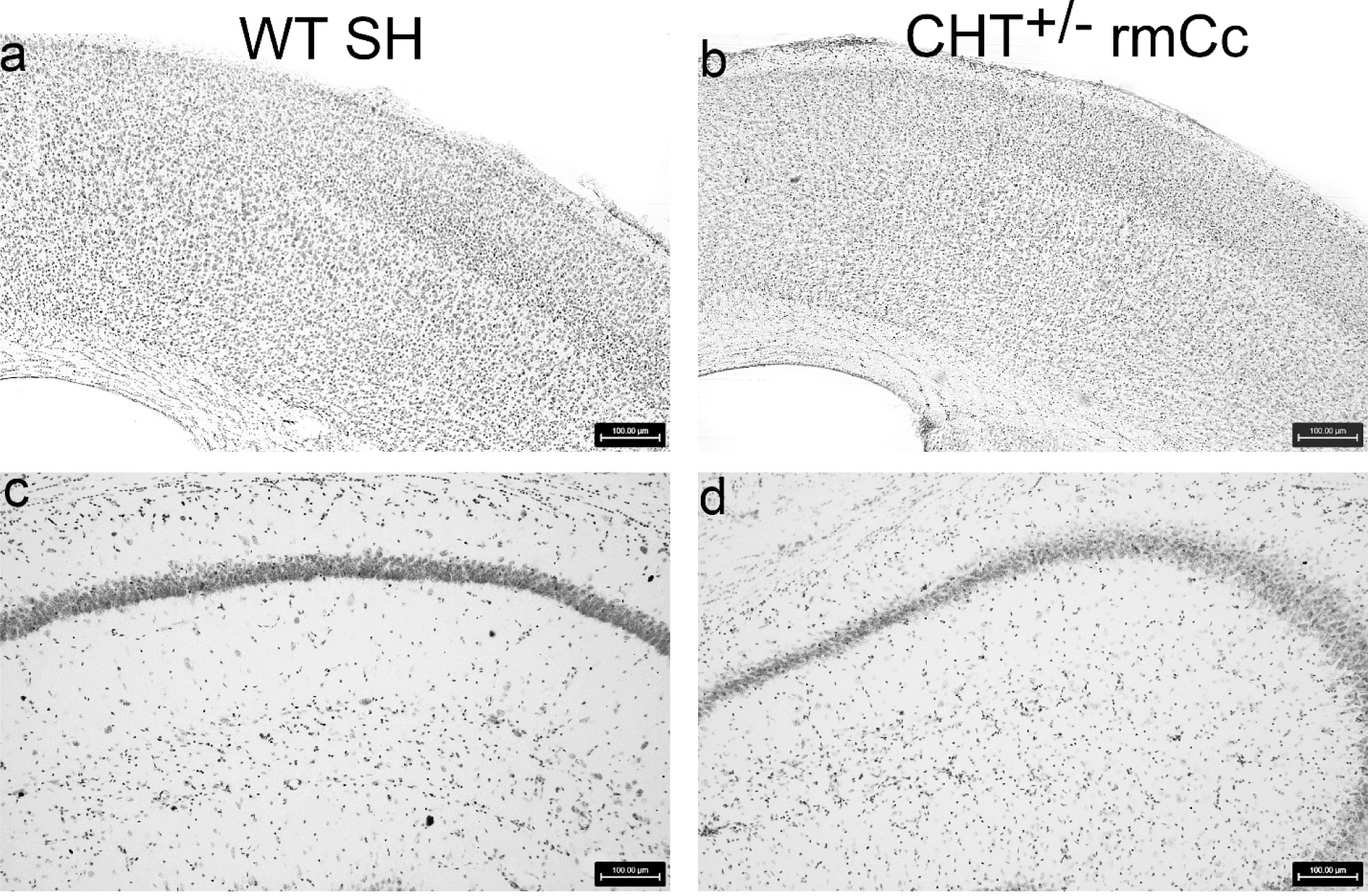
Nissl-stained agranular motor and granular somatosensory motor (a,b), and dorsal hippocampus (c,d), of a WT SH and a CHT^+/-^ rmCc mouse (100 µm scale shown in <u>a-d</u>). As was expected from prior studies using a comparable method to generate rmCc (Kane et al., 2012), the relatively milder rmCc method employed in the present experiments did not yield evidence of apparent macroanatomical damage or gliosis in the brains of wild type of CHT^+/-^ mice.

## Discussion

We hypothesized that CHT^+/-^ mice are relatively more vulnerable than WT mice to the cognitive and neuronal effects of rmCc. The impact of our rmCc procedure was relatively limited as indicated by the absence of effects on attentional performance of WT mice, the macroanatomical integrity of the brains of all groups, and the density of cortical synapses as indicated by synaptophysin levels. In CHT^+/-^ mice, rmCc lastingly impaired the attentional performance. This effect was associated with and, as will be discussed below, likely mediated by, a nearly complete loss of the capacity of cholinergic synapses to import choline via the CHT.

Cholinergic neurotransmission in CHT^+/-^ mice is unaltered at baseline (Ferguson et al., 2004) but exhibits an attenuated capacity for sustaining elevated levels of activity, such as seen during SAT performance (Paolone, Mallory, et al., 2013). In CHT^+/-^ mice, and in the absence of stimulation of cholinergic neurons, the overall reduction of CHT protein solely reflects a decrease in intracellular CHT density, with synaptosomal plasma membrane CHT density being preserved at WT level (Ferguson et al., 2004; Ferguson et al., 2003; Parikh et al., 2013). In the absence of stimulation, this situation supports WT levels of CHT-mediated choline transport and ACh release in CHT^+/-^ mice. In contrast, stimulation of cholinergic neurons, such as during SAT performance, reveals the attenuated outward mobilization of intracellular CHTs and thus the attenuated capacity for elevated cholinergic neurotransmission (Ferguson et al., 2003; Parikh et al., 2013).

Cholinergic systems mediate two essential aspects of SAT performance. Fast, phasic or transient cholinergic signaling is necessary for scoring hits (Gritton et al., 2016; Howe et al., 2017; Sarter, Lustig, Berry, et al., 2016) while a slower, neuromodulatory component mediates levels of attentional control, including task representation and maintenance, and performance stabilization and recovery following performance challenges (Kim et al., 2017a; Sarter & Lustig, 2019; Sarter, Lustig, Blakely, et al., 2016; Sarter, Lustig, & Taylor, 2012; Sarter & Paolone, 2011; St Peters, Demeter, et al., 2011). In CHT^+/-^ mice, and in the absence of additional performance or cholinergic challenges, SAT performance remains sufficiently supported by residual levels of cholinergic activity (Paolone, Mallory, et al., 2013; Parikh et al., 2013), likely reflecting that cholinergic neurons remain the capacity for generating phasic ACh release events, and that well-practiced SAT performance only minimally taxes attentional control. Following rmCC, however, CHT capacity in CHT^+/-^ mice was nearly completely attenuated and thus likely interfered also with the generation of cholinergic transients. Consistent with our prior demonstration of effects of optogenetic silencing of cholinergic transients on SAT performance (Gritton et al., 2016), rmCc-induced near-silencing of CHT capacity in CHT^+/-^ mice was associated with a permanent suppression of hit rates in these mice.

rmCc reduced SAT-activated choline uptake in WT mice by about 50% relative to WT SH rats. In CHT^+/-^ mice, rmCc also reduced choline uptake by about 50% relative to their sham-treated counterparts. However, this effect in CHT^+/-^ mice amounted to an approximately 75% decrease in CHT capacity relative to sham-treated WT mice and, importantly, in CHT^+/-^ mice, rmCC nearly completely attenuated CHT-mediated choline transport (with residual Vmax scores around 2 pmol/mg protein/5 min; Fig. 3b).

For the further interpretation of these findings, it is important to note that choline uptake was measured in mice performing the SAT. As was frequently demonstrated, behavioral performance can alter synaptosomal choline uptake (Decker, Pelleymounter, & Gallagher, 1988; Messier, Durkin, Mrabet, & Destrade, 1990). Indeed, in WT rats and mice, SAT performance upregulates cortical choline transport, mediated via an increased rate of translocation of CHTs from intracellular sites to plasma membrane (Apparsundaram et al., 2005; Parikh et al., 2013). In CHT^+/-^ mice, however, SAT performance did not elevate plasma membrane CHT density (Parikh et al., 2013), and thus SAT performance was unlikely to elevate choline uptake in CHT^+/-^ SH mice. As rmCc exerted similar degrees of reduction of SAT-associated choline transport when compared with sham-operated controls of the same genotype, it is possible that similar cellular mechanisms mediated the effects of rmCc in both genotypes. These mechanisms remain unknown but neither involved the loss of CHTs nor, as suggested by the unchanged synaptophysin protein levels, loss of cortical synapses in general. While our data do not support a gross change in overall synaptic density, more selective measures of cholinergic terminals independent of acetylcholine homeostasis must be implemented to rule out a frank loss of these elements.

Although the present results did not suggest that rmCc produced a fundamentally different effect in WT versus CHT^+/-^ mice, the absolute net effect in CHT^+/-^ mice appears essential for understanding their lasting impairments in SAT performance. As shown before, the residual cholinergic capacity of CHT^+/-^ mice is sufficient to support (unchallenged) SAT performance (Paolone, Mallory, et al., 2013), and the present results confirmed this finding. Moreover, after rmCc, WT likewise remain the capacity for such performance, consistent with SAT-associated choline uptake levels that matched those seen in CHT^+/-^ SH mice. However, in CHT^+/-^ mice, rmCC reduced choline uptake to levels that likely were too low to support residual cholinergic mediation of SAT performance. Together, these findings suggest an additive effect of genotype and rmCC on choline transport which, in CHT^+/-^ mice, yielded a robust and lasting cognitive deficit.

Our ongoing studies have indicated the possibility that the effects of rmCc in CHT^+/-^ mice include lasting increases in the activity of pro-neuroinflammatory brain cytokines (Koshy Cherian, Tronson, Parikh, Blakely, & Sarter, 2015). If confirmed, these findings would suggest escalating detrimental interactions between a relatively low capacity for cholinergic neurotransmission in CHT^+/-^ mice *per se*, and an exaggerated neuroinflammatory response to rmCc which may further suppress CHT function (see also Zhu, Blakely, & Hewlett, 2006),

The main results from this experiment predict that in subjects with a relatively low capacity for sustaining elevated levels of cholinergic neurotransmission, even relatively mild concussions will yield severe and lasting cognitive impairments. Thus, not only is the forebrain cholinergic system exquisitely vulnerable to the effects of brain injury (Dixon et al., 1996; Dixon et al., 1994; Dixon, Ma, & Marion, 1997; Gorman, Fu, Hovda, Murray, & Traystman, 1996; Salmond et al., 2005), but a vulnerable cholinergic system represents a major risk factor for developing severe cognitive decline after even mild impacts (see also Conner et al., 2005; Field, Gossen, & Cunningham, 2012). Our research in humans heterozygous for a low capacity CHT version, present in about 10% of the population, indicated distinct attentional vulnerabilities, an associated failure to activate right prefrontal regions, and vulnerability for depression (Berry et al., 2015; Berry et al., 2014; English et al., 2009; Hahn et al., 2008; Sarter, Lustig, Blakely, et al., 2016). The present data predict that these subjects are also more vulnerable to the effects of rmCc. Likewise, Parkinsonian fallers, who in addition to dopamine losses also exhibit cholinergic losses (Bohnen et al., 2009), and consequently attentional impairments (Kim et al., 2017a, 2017b), would be predicted to suffer from relatively severe consequences of even very mild concussions, thereby possibly further accelerating the progress of the disease (e.g., Perry et al., 2016). Given the essential role of CHT capacity for cholinergic neurotransmission, CHT^+/-^ mice appear to be a useful model for future research to study the inflammatory mediators of such cholinergic vulnerabilities and to identify potential therapies for limiting the severe effects of mild impacts in vulnerable humans.

Author Contributions
A.K.C., N.C.T. and M.S. designed research. A.K.C., N.C.T. and V.P. performed research. A.K. analyzed the data. R.D.B. contributed reagents/analytical tools. A.K.C., N.C.T, R.D.B. and M.S. wrote the paper.

## Acknowledgements

This research was supported by PHS grant MH086530 (MS) and the Dystonia Medical Research Foundation (RDB). We thank Marc Bradshaw (University of Michigan) for designing, building and estimating the force of the impact device used in this study. A preprint of this paper was posted online on bioRxiv (doi: http://dx.doi.org/10.1101/521484).

## Notes

Competing Financial Interests: The authors declare no competing financial interest.

